# Highly efficient intercellular spreading of protein misfolding mediated by viral ligand - receptor interactions

**DOI:** 10.1101/2020.06.26.173070

**Authors:** Shu Liu, Andre Hossinger, Annika Hornberger, Oleksandra Buravlova, Stephan Müller, Stefan F. Lichtenthaler, Manuela Neumann, Philip Denner, Ina M. Vorberg

## Abstract

Pathological protein aggregates associated with neurodegenerative diseases have the ability to transmit to unaffected cells, thereby templating their own aberrant conformation onto soluble proteins of the same kind. Proteopathic seeds can be released into the extracellular space, secreted in association with extracellular vesicles (EV) or exchanged by direct cell-to-cell contact. The extent to which each of these pathways contributes to the prion-like spreading of protein misfolding is unclear. Exchange of cellular cargo by both direct cell-to-cell contact as well as via EV depends on receptor-ligand interactions and subsequent release of cargo into the cytosol. We hypothesized that enabling these interactions through viral ligands enhances the aggregate-inducing capacity of EV-associated proteopathic seeds. Using different cellular models propagating model prion-like protein aggregates, mouse-adapted prions or pathogenic Tau aggregates, we demonstrate that vesicular stomatitis virus glycoprotein and SARS-CoV-2 spike S increase protein aggregate induction by direct cell-to-cell contact or via viral glycoprotein-decorated EV. Thus, receptor-ligand interactions are major determinants of intercellular aggregate dissemination. Further, our data raise the intriguing possibility that acute or latent viral infections contribute to proteopathic seed spreading by facilitating intercellular cargo transfer.

**HIGHLIGHTS:**

- Different types of proteopathic seeds are secreted in association with extracellular vesicles
- Receptor-ligand interactions are important drivers of direct cell-to-cell and extracellular vesicle-mediated spreading of protein misfolding
- Viral glycoproteins mediating attachment and membrane fusion strongly enhance aggregate inducing capacity in recipient cells

**GRAPHICAL ABSTRACT:** 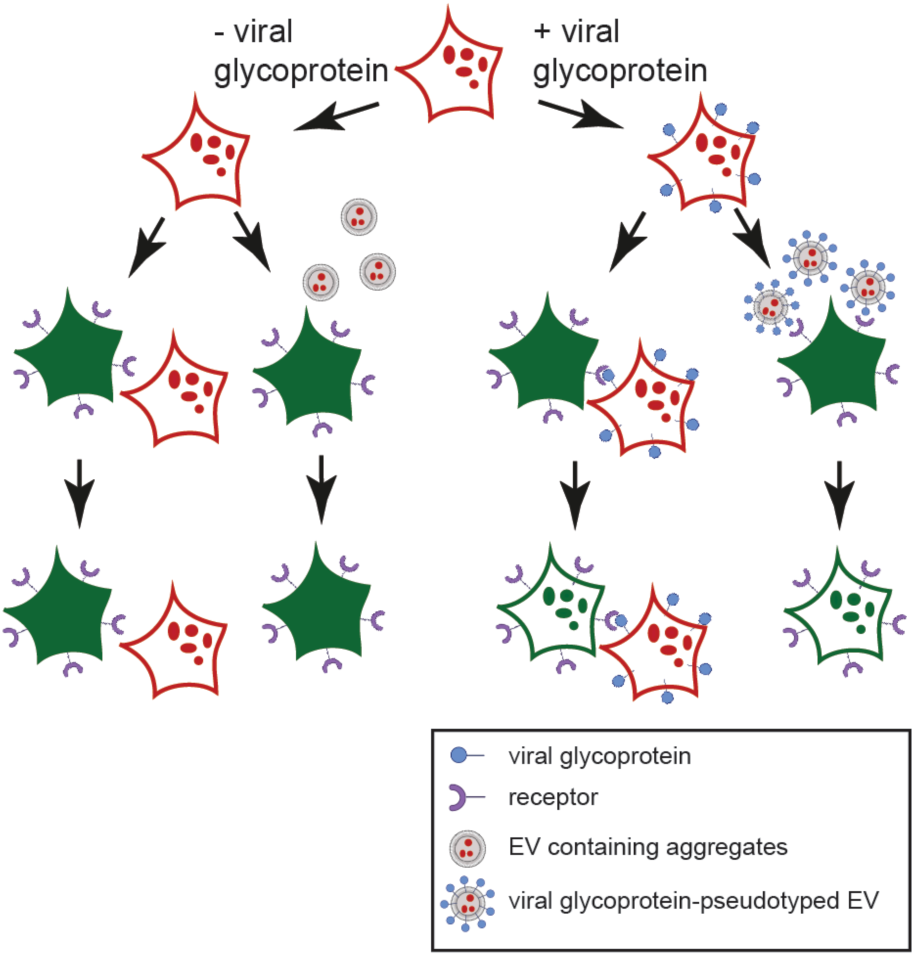

## INTRODUCTION

Aberrant folding and aggregation of host-encoded proteins into ordered protein assemblies is a pathological hallmark of neurodegenerative diseases such as prion diseases, Alzheimer’s disease (AD) and Parkinson’s disease (PD). Disease-associated protein deposition usually starts locally, subsequently spreading stereotypically to other brain regions ^1^. AD, the most prevalent neurodegenerative disease, is associated with the extracellular accumulation of Aβ amyloid and intracellular inclusion of misfolded microtubule-binding protein Tau as in neurofibrillary tangles ^2, 3^. The formation of Tau aggregates is also a hallmark of other neurodegenerative diseases, collectively known as tauopathies ^4^. In PD, normally soluble α-synuclein is deposited into aberrant neuronal inclusions ^5^. Transmissible spongiform encephalopathies or prion diseases constitute a special group of neurodegenerative diseases affecting humans and other mammals ^6^. Prion diseases can occur sporadically, be caused by mutations in the prion protein gene *PRNP* or can be acquired by infection or iatrogenic transmission ^7^.

Pathogenic protein aggregates form in a time-limiting nucleation dependent process in which soluble aggregation-prone proteins form oligomers that subsequently grow into highly ordered, beta-sheet rich fibrils ^8^. *In vitro* assays demonstrate that the lag phase required for seed formation drastically shortens in the presence of pre-formed polymers that act as seeds ^9^. Interestingly, such proteopathic seeds not only recruit monomeric protein in the affected cell, but also in unaffected cells upon intercellular transmission, a process that likely underlies the often observed stereotypical spreading of protein misfolding in neurodegenerative diseases ^1^. In fact, intercellular transmission of proteopathic seeds is regarded as a common mechanism of neurodegenerative diseases ^10^. The precise mechanism of intercellular aggregate transfer and induction of new aggregates is unclear, but appears to involve release of free protein aggregates, direct cell-to-cell contact by cytonemes such as tunneling nanotubes (TNTs) ^11-13^, or release of extracellular vesicles (EV) containing small proteopathic seeds ^14^. The extent to which these three routes contribute to the spreading of protein misfolding remains unclear.

EV are nanosized communication vesicles secreted by all cells under physiological and pathological conditions ^15^. EV differ in their cellular origin, with some budding directly from the cell surface (so-called microvesicles), and others being secreted when multivesicular bodies of endosomal origin fuse with the plasma membrane and release exosomes into the extracellular space. Secreted vesicles are now generally referred to as EV due to substantial overlap of microvesicles and exosomes in terms of size, surface markers and function ^16^. Soluble and aggregated isoforms of a diverse number of neurodegenerative diseases have been found secreted by neurons and other cells either as free proteins or in association with EV ^17-21^. While EV containing proteopathic seeds have been shown to seed protein aggregation *in vitro* and *in vivo*, the efficiency of protein aggregate transfer and subsequent seeding through this route is unclear. Only a small fraction of soluble or aggregated proteins are released associated with EV, while the vast majority is freely secreted. For example, less than 1% of total secreted Aβ associated with EV ^17^, and only 3% of total secreted α-synuclein was found in the EV fraction ^22^. Rat cortical neurons have been shown to secrete Tau, but again very little (3%) was associated with EV ^19^. Importantly, EV fractions purified from conditioned medium of N2a cells overexpressing an aggregated Tau mutant induced Tau aggregation in less than 0,1% of recipient cells, arguing that EV-mediated aggregate induction can be rather inefficient ^19^.

EV isolated from different donor cells exhibit a marked cell tropism ^23, 24^. Independent of their uptake mechanisms, EV must merge with cellular membranes to release their cargo into the cytosol. EV docking onto target cells and subsequent uptake are selective processes that require specific membrane interactions ^24^. Consequently, receptor-ligand interactions between EV and recipient cells will likely modulate the spreading behavior of proteopathic seed cargo. While integrins and proteoglycans have been identified that adhere EV to target cells, most receptor-ligand pairs that underlie these targeted interactions are so far unknown ^24, 25^.

We reasoned that the poor aggregate inducing activity of some seed-containing EV could be due to the lack of specific ligands required for host receptor interactions and/or subsequent fusion processes. Membrane contact and fusion can be facilitated by viral glycoproteins that allow viruses to adhere to and penetrate into their target cells. The contact with specific receptors on apposed membranes leads to conformational transitions in the viral glycoproteins, thereby bringing the two membranes in close proximity and enforcing bilayer merger. Viral glycoproteins are routinely used to pseudotype genetically engineered viral vectors for efficient cargo delivery.

The vesicular stomatitis virus glycoprotein VSV-G is the sole surface protein of this virus belonging to the rhabdovirus genus that mediates the binding to the LDL receptors present on most cell types ^26^. A conformational change in VSV-G subsequently results in fusion of host and viral lipid bilayers ^27^. Ectopically expressed VSV-G is not only found on the cell surface but also decorates EV in the absence of other viral constituents ^28^. It has been recently demonstrated that VSV-G enhances EV-mediated cargo delivery to recipient cells ^28, 29^. We here expressed VSV-G in three different cellular models that propagate different protein aggregates. Coculture of VSV-G-expressing donor cells with recipient cells strongly increased protein aggregate induction in the latter. Further, expression of VSV-G also promoted the secretion of VSV-G-coated EV with drastically enhanced aggregate-inducing capacity in recipient cells. Intriguingly, interactions between SARS-CoV-2 spike S protein and its receptor ACE2 similarly contributed to the spreading of cytosolic prions and Tau aggregates. Thus, efficient intercellular proteopathic seed transfer is strongly controlled by receptor-ligand interactions. Further, our data raise the intriguing possibility that viral glycoproteins, expressed during acute or chronic infection, could facilitate the spreading of protein misfolding *in vivo*.

## Results

### Expression of viral glycoprotein VSV-G drastically increases cell-to-cell spreading of cytosolic prions

To study the intercellular dissemination and propagation of proteinaceous seeds, we have implemented cellular models that rely on the ectopic expression of a yeast prion domain in the cytosol of mammalian cells. The *Saccharomyces cerevisiae* Sup35 translation termination factor is the prototype prion protein of lower eukaryotes. Its classification as a prion protein is based on the fact that it can exist in a functional soluble isoform and as an inactive, cross-beta sheet polymer that self-replicates by imprinting its conformation onto soluble protein of the same kind ^30^. When expressed in the mammalian cytosol, the NM prion domain of Sup35 exists in a soluble state and is non-functional ^31^. Exposure of cells to highly ordered protein fibrils composed of recombinant NM leads to the aggregation of the ectopically expressed NM, thereby inducing self-replicating NM prions that are heritable by progeny ^31^. NM prions are also transmissible to naïve bystander cells by direct cell-to-cell contact ^32^ and EV ^33^.

Interestingly, we found that donor cell populations drastically differ in their NM aggregate-inducing activity, an effect that also depended on the recipient cell line ^33, 34^. For example, coculture of HEK cells propagating HA-epitope tagged NM prions (NM-HA^agg^) only poorly induced NM-GFP prions (NM-GFP^agg^) in recipients (**suppl. figure 1**). Likewise, EV secreted by HEK NM-HA^agg^ cells contained substantial amounts of aggregated NM ^34^, yet were ineffective at inducing NM-GFP aggregation in recipient cells (**suppl. figure 1**). We hypothesized that one reason for the poor NM aggregate induction in some recipient cell populations could be inefficient membrane contact and fusion of either EV with target cells or donor and recipient cells. Since vesicular stomatitis virus glycoprotein VSV-G binds to the broadly expressed LDL receptor and therefore increases cell-to-cell membrane contact ^26^, we assessed whether the expression of VSV-G facilitates protein aggregate transmission. We first tested if ectopic VSV-G expression increased cell-to-cell spreading of NM prions from donor to recipient N2a or HEK cells (**Fig. 1a**). As donors, we chose HEK NM-HA^agg^ cells ^34^ and N2a NM-HA^agg^ clone 2E ^32^, which have low NM aggregate induction rates when cocultured with recipient cells. Donor cells were transfected with a plasmid coding for VSV-G and cells were subsequently cocultured with recipient N2a or HEK cell lines stably expressing soluble NM-GFP (NM-GFP^sol^) (**Fig. 1b; suppl. figure 2a**). Independent of donor and recipient cell lines, expression of VSV-G by donor cells drastically increased the percentage of recipient cells with induced NM-GFP aggregates up to 50-fold upon 16 hours of coculture (**Fig. 1c-f**). In a few instances, we observed syncytia formation between donor and recipient cells, possibly induced by the fusogenic activity of VSV-G (**suppl. figure 2b**). No NM-GFP aggregate formation was observed when donor cells expressed only soluble NM-HA (**suppl. figure 2c, d**). The prion-inducing activity of donor cells expressing VSV-G strongly depended on the fusogenic activity of the viral protein, as donor cells expressing a non-fusogenic VSV-G mutant exhibited a drastically reduced NM-GFP aggregate inducing activity in recipient cells (**Fig. 1g, h**).

**Fig. 1.**
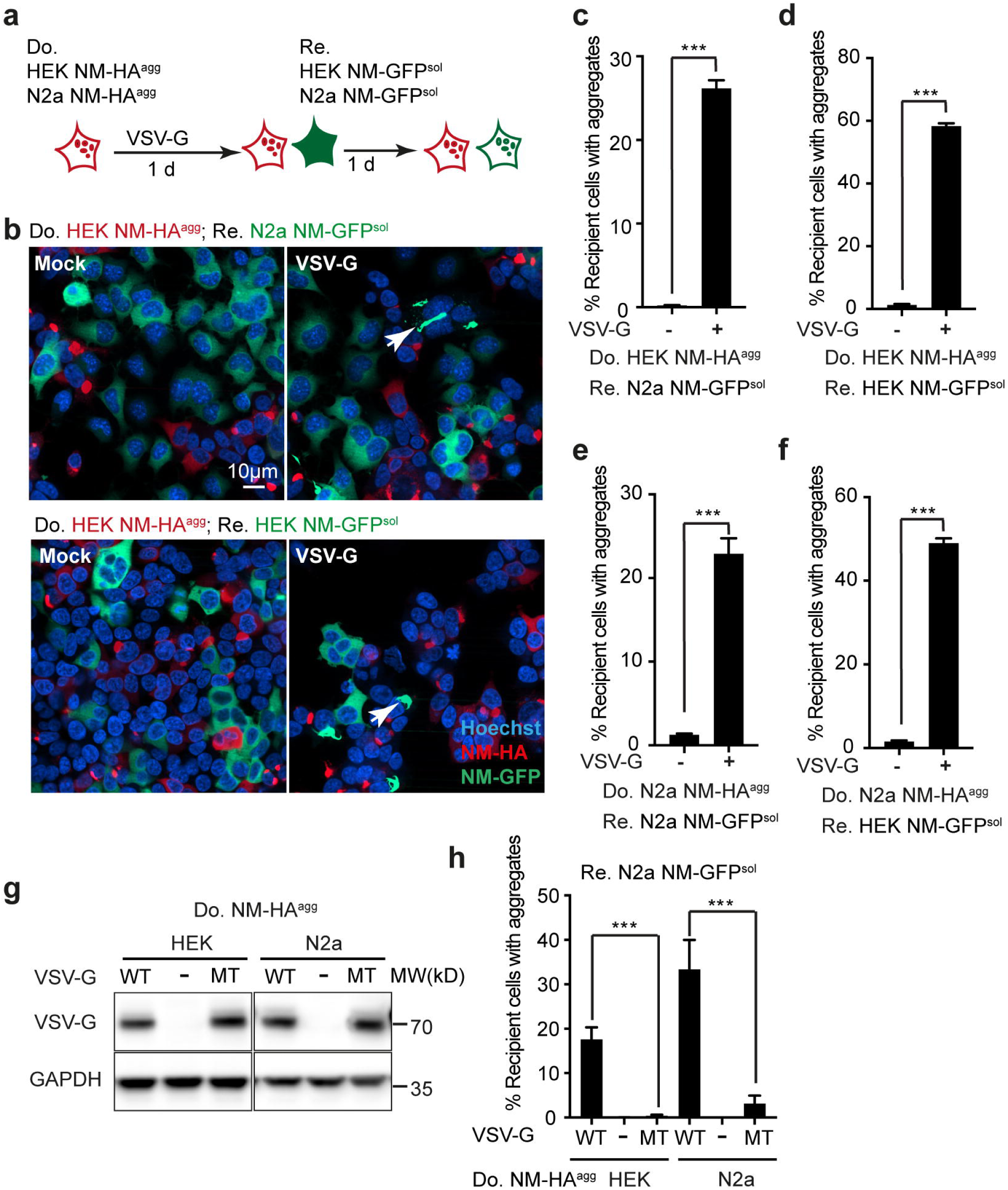
VSV-G expression by donor cells increases induction of Sup35 NM prions in cocultured recipient cells. **a**. Experimental design. HEK NM-HA^agg^ cell clone (red) and N2a NM-HA^agg^ clone (red) with poor aggregate inducing activity in recipients were chosen as donors. Donor cells were transiently transfected with plasmid coding for VSV-G or Mock transfected and subsequently cocultured with recipient HEK or N2a cells expressing NM-GFP^sol^ (green). **b**. Representative confocal microscopy images of transfected HEK NM-HA^agg^ donor cells cocultured with N2a NM-GFP^sol^ (top panel) or HEK NM-GFP^sol^ cells (bottom panel). Do.: Donor; Re.: Recipient. NM-HA was stained using anti-HA antibodies (red). Nuclei were stained with Hoechst. Scale bar: 10 µm. **c**.**-f**. Quantitative analysis of the percentage of recipient cells with induced NM-GFP aggregates (NM-GFP^agg^). Statistical analysis was performed using student’s unpaired t-test (n=6, ***, *p*<0.001). **g**. Donor HEK NM-HA^agg^ or N2a NM-HA^agg^ cells were transfected with empty vector or vectors coding for active (WT) or a non-fusogenic VSV-G^W72A, Y73A^ variant (MT). Western blot shows VSV-G expression of donor cells. GAPDH served as loading control. **h**. Mock transfected (-), VSV-G (WT) or VSV-G^W72A, Y73A^ mutant (MT) transfected donor cells were cocultured with recipient N2a NM-GFP^sol^ cells. NM-GFP aggregate induction was determined one day post coculture (n=6, ***, *p*<0.001; one-way ANOVA).

### VSV-G mediates efficient vesicular dissemination of cytosolic NM prions

VSV-G is a type III fusogen present in an inactive pre-fusion state at neutral pH that becomes activated by low pH in the endolysosomal system ^35^. The occasional syncytia formation argued that at least some of the aggregate induction events could be due to extracellular VSV-G activation and subsequent cell fusion rather than contact and fusion at local cell contacts, such as cytonemes. We thus tested the effect of VSV-G expression on cell-free NM-GFP aggregate induction. EV were isolated from N2a or HEK donor cells transfected with empty vector or VSV-G coding vector. EV were subsequently added to recipient N2a or HEK cells NM-GFP^sol^ cells (**Fig. 2a**). EV fractions isolated from donor populations contained VSV-G, NM-HA and EV marker proteins Flotillin1, Hsp70/72, ALIX and GAPDH (**Fig. 2b**). The presence of VSV-G on EV isolated from donor N2a or HEK NM-HA^agg^ cells strongly increased intercellular aggregate induction when EV were added to HEK cells expressing NM-GFP^sol^ (**Fig. 2c**). Sonication of EV fractions prior to their addition to cells strongly reduced aggregate induction, demonstrating that intact EV were required for efficient prion spreading (**Fig. 2c**). Furthermore, expression of VSV-G proved non-toxic to donor cells in our experimental setup (**Fig. 2d**). Thus, the increased aggregate induction efficiency was unlikely due to free naked NM-HA aggregates released by dying cells.

**Fig. 2.**
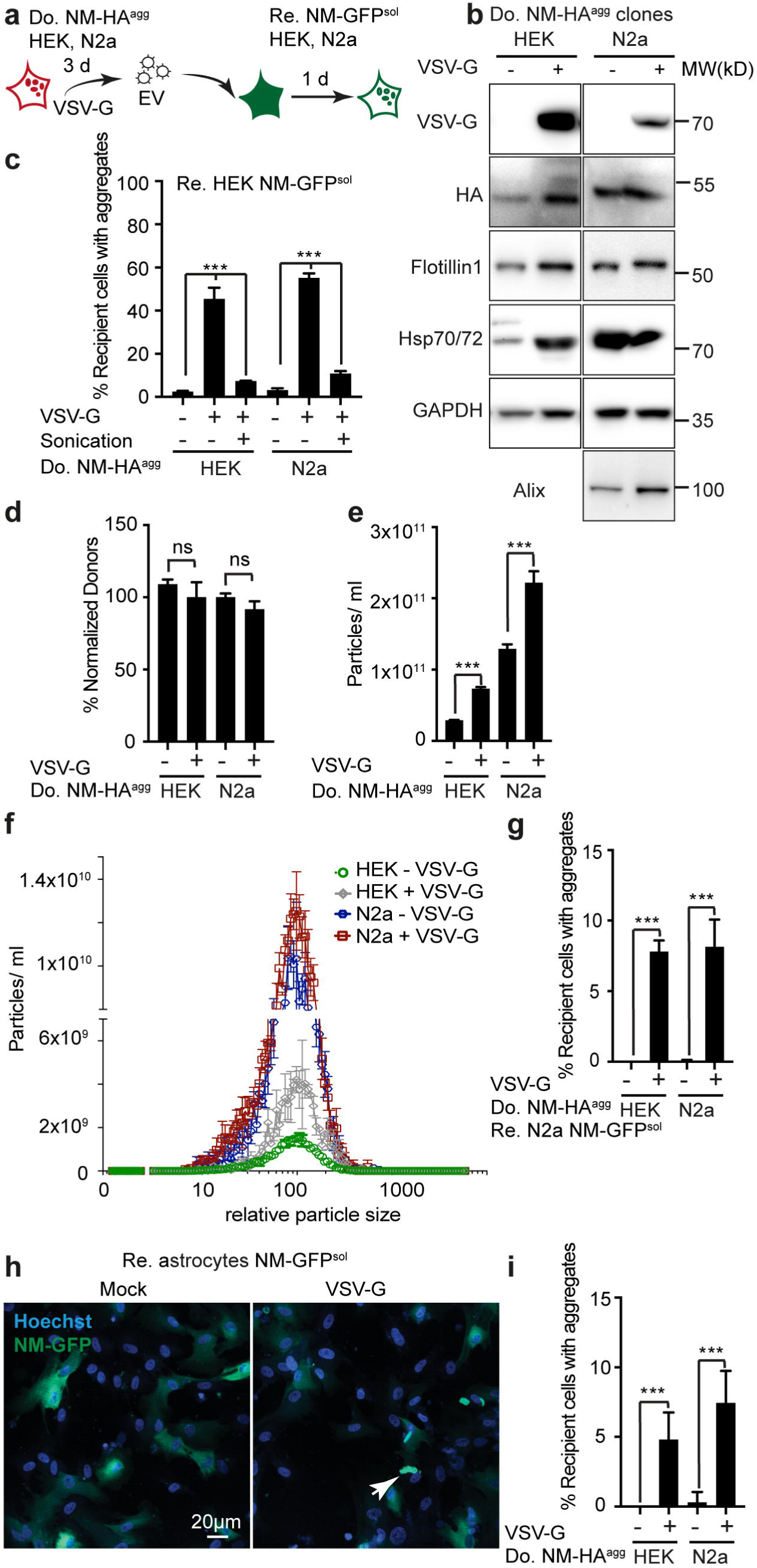
Pseudotyping extracellular vesicles with VSV-G drastically increases their intercellular NM aggregate inducing efficiency. **a**. Experimental workflow. HEK or N2a cell clones propagating aggregated NM-HA (NM-HA^agg^) were transfected with VSV-G coding plasmid or empty vector. EV from donor clones were isolated from conditioned medium and added to recipient HEK or N2a cells expressing NM-GFP^sol^. **b**. EV were assessed for the presence of EV markers and NM-HA by Western blot. **c**. Percentage of recipient HEK NM-GFP cells with induced NM-GFP aggregates upon exposure to EV produced by Mock- or VSV-G-transfected donor HEK or N2a NM-HA^agg^ cells. Sonication was used to destroy EV (n=6, ***, *p*<0.001; one-way ANOVA). **d**. Expression of VSV-G by donor cells is non-toxic. Total cell numbers of donor cells transfected with empty vector (Mock) or vector coding for VSV-G were assessed three days post transfection. Mock control was used for normalization. Statistical analysis was performed using student’s unpaired t-test (n=6, ns: non-significant). **e**. Particle numbers of EV in conditioned medium from either VSV-G-transfected or Mock transfected donor N2a NM-HA^agg^ or HEK NM-HA^agg^ were determined three days post transfection using the ZetaView nanoparticle tracking device. Student’s unpaired t-test (n=3, ***, *p*<0.001). **f**. Size distribution graph of different EV revealed no effect of VSV-G on size of EV. **g**. EV preparations adjusted for comparable EV numbers were added to recipient N2a NM-GFP^sol^ cells. Cells fixed 16 h post EV addition were analyzed for the percentage of cells with induced NM-GFP aggregates. Student’s unpaired t-test (n=6, ***, *p*<0.001). **h**. Human astrocytes expressing NM-GFP^sol^ were exposed to VSV-G pseudotyped or non-pseudotyped EV and subsequently assessed for NM-GFP aggregates 1 day later. Nuclei were stained with Hoechst. Scale bar: 20 µm. **i**. Percentage of astrocytes expressing NM-GFP with induced NM-GFP aggregates upon exposure to EV from either HEK or N2a cells. Statistical analysis was performed using student’s unpaired t-test (n=6, ***, *p*<0.001).

The effect of VSV-G expression on prion induction could either be due to enhanced production of EV, enhanced membrane docking and fusion of EV and recipient cells, or both. Consistent with previous findings ^29^, we observed an approximately two-fold increase in EV release upon transient expression of VSV-G in donor cell clones (**Fig. 2e**). VSV-G pseudotyping did not change the size distribution of released EV (**Fig. 2f)**. When adjusted for particle numbers, still an increase in intercellular aggregate induction was observed when EV were pseudotyped with VSV-G (**Fig. 2g**). Pseudotyped EV secreted by donor cell lines also induced NM-GFP aggregation in primary human astrocytes ectopically expressing soluble NM-GFP, demonstrating that VSV-G mediated aggregate induction was not restricted to permanent cell cultures (**Fig. 2h, i)**. We conclude that VSV-G expressed by donor cells can strongly increase intercellular NM prion induction by mediating efficient contact and fusion of donor and recipient cell membranes as well as of EV and target cells.

### Enhanced intercellular transmission of Tau aggregation upon VSV-G expression

Accumulating evidence suggests that seeding-competent Tau species can spread from cell-to-cell by EV ^19^. To assess the effect of receptor-ligand interactions on intercellular Tau aggregate induction, we employed a previously published Tau cell model ^36^. We engineered HEK cells to stably express the aggregation competent Tau repeat-domain-spanning amino acid residues 244-372 (carrying the two-point mutations P301L/V337M and fused to GFP - hereafter termed Tau-GFP) and exposed them to homogenates extracted from affected brain regions from patients with Alzheimer’s disease (AD), cortical basal degeneration (CBD), progressive supranuclear palsy (PSP) or frontotemporal lobar degeneration with Tau pathology (FTLD-Tau). All patient brain homogenates contained aggregated Tau, as revealed by pronase digestion (**suppl. figure 3a**). Upon limiting dilution cloning, we established HEK cell clones Tau-GFP^AD^, Tau-GFP^FTLD^, Tau-GFP^PSP^ and Tau-GFP^CBD^ stably producing Tau aggregates (**Fig. 3a**). Sedimentation assays and pronase treatment demonstrated the presence of aggregated Tau-GFP in all cell clones exposed to patient brain homogenate, but not in control cells exposed to control brain homogenate (**Fig. 3b, suppl. figure 3b**).

**Fig. 3.**
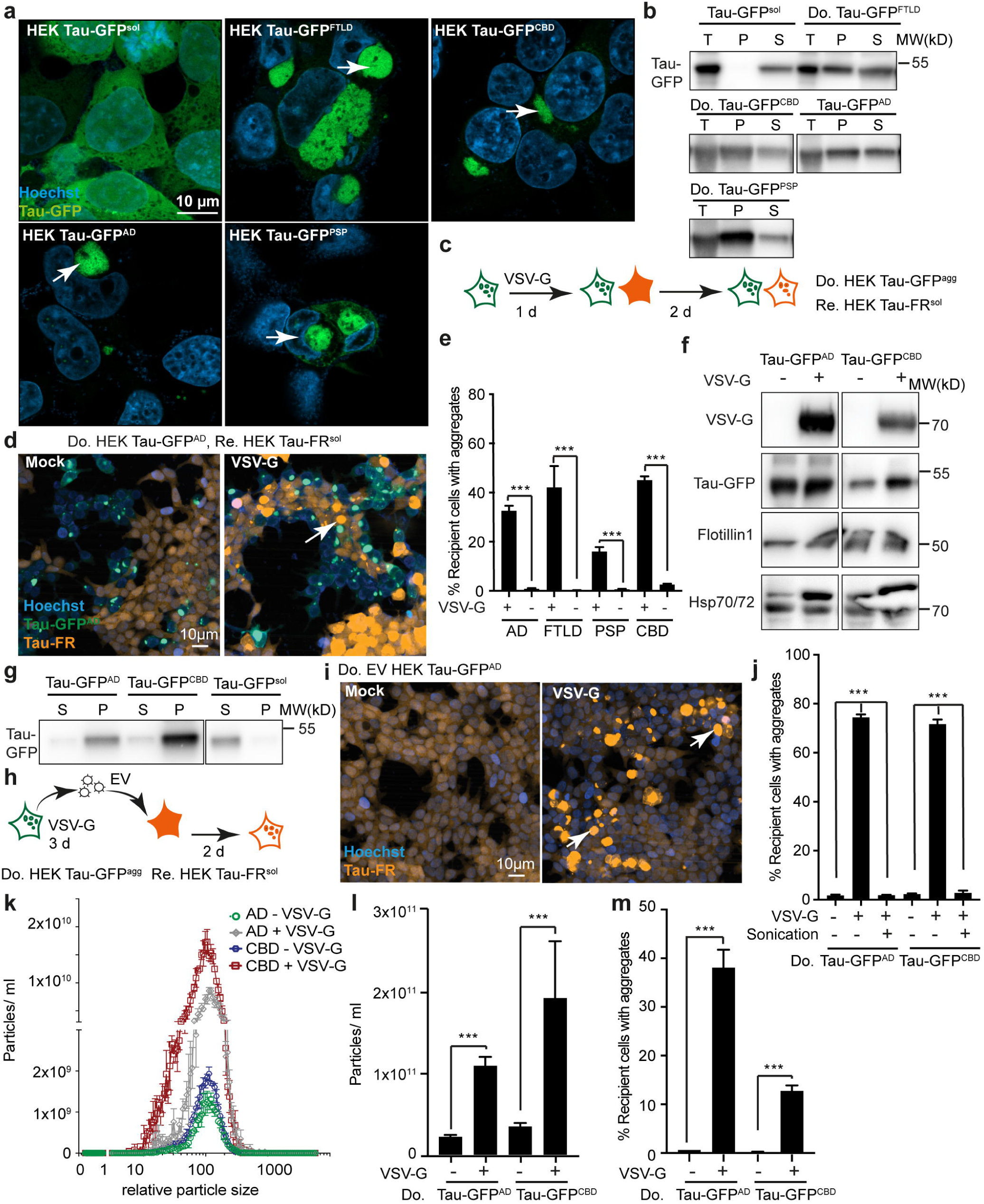
VSV-G expression enhances cell-to-cell and cell-free induction of Tau aggregates. **a**. Immunofluorescence staining of the donor HEK Tau-GFP population and HEK clones propagating aggregated Tau-GFP. A HEK cell clone expressing soluble Tau-GFP (HEK Tau-GFP^sol^) was exposed to final 1 % post mortem brain homogenates from frozen postmortem brain tissue from tauopathy patients (AD: Alzheimer’s disease; CBD: Corticobasal degeneration; FTLD-tau: Frontotemporal lobar degeneration with tau-pathology; PSP: Progressive supranuclear palsy). Limiting dilution cloning was performed to isolate single cell clones that stably form visible Tau-GFP foci (Tau-GFP^agg^, also termed Tau-GFP^AD^, Tau-GFP^FTLD^, Tau-GFP^PSP^, Tau-GFP^CBD^, depending on the brain homogenate used for induction). Nuclei were stained with Hoechst. Arrowheads indicate Tau-GFP foci in different cell clones. **b**. Sedimentation assay of cell lysates from uninduced HEK Tau-GFP^sol^ cells and individual clones propagating Tau-GFP aggregates. T: Total cell lysate; P: insoluble pellet fraction; S: soluble supernatant fraction. **c**. Workflow of the experiment. HEK Tau-GFP^agg^ clones were transfected with vector coding for VSV-G or the empty vector. Recipient HEK cells stably expressing soluble Tau-FusionRed (HEK Tau-FR^sol^) were cocultured with transfected donor cells. **d**. Donor clone HEK Tau-GFP^AD^ was Mock transfected or transfected with VSV-G plasmid and subsequently cocultured with recipient HEK Tau-FR^sol^. Foci induction was monitored 2 days post coculture. Arrow indicates induced Tau-FR foci. **e**. Percentage of recipient cells with induced Tau-FR foci following coculture with donors (n=6, ***, *p*<0.001; one-way ANOVA). **f**. Presence of VSV-G, Tau-GFP and EV markers Flotillin1, Hsp70/72 in EV fractions of donor cell lines. **g**. Sedimentation assay using EV isolated from HEK Tau-GFP^AD^, HEK Tau-GFP^CBD^ and uninduced HEK Tau-GFP^sol^ cells demonstrates that Tau-GFP packaged in EV is predominately in an insoluble state. Tau-GFP was detected using anti-Tau antibody (ab64193). **h**. Experimental overview. Different HEK Tau-GFP^agg^ cell clones were transiently transfected with a plasmid coding for VSV-G or were Mock transfected. Three days later, EV were harvested and added to recipient HEK Tau-FR^sol^ cells. **i**. Exposure of recipient HEK Tau-FR^sol^ cells to EV isolated from conditioned medium of HEK Tau-GFP^AD^ donors with / without VSV-G expression. **j**. Percentage of recipient cells with induced Tau-FR aggregates following VSV-G pseudotyped exosome addition. Sonication was used to destroy EV (n=6, ***, *p*<0.001; one-way ANOVA). **k**. Size distribution graph of different EV is shown (analyzed by Zetaview). **l**. Particle numbers released upon transfection with vector coding for VSV-G. HEK Tau-GFP^AD^ and HEK Tau-GFP^CBD^ cell clones were Mock transfected or transfected with VSV-G coding vector. Particles released into the medium were determined 3 d post transfection. **m**. Isolated EV were adjusted to comparable particle numbers and added to recipient HEK-FR^sol^ cells. HEK Tau-FR cells with induced TauFR^agg^ were determined 2 d post EV addition. Statistical analysis was performed using one-way ANOVA (n=6, ***, *p*<0.001).

Generated donor cell clones propagating Tau aggregates were either transfected with empty vector or vector coding for VSV-G. Donor clones were subsequently cocultured with recipient cells expressing soluble Tau-FusionRed (Tau-FR^sol^) (**Fig. 3c**). Coculture with Mock transfected donor cells resulted in poor induction of Tau-FR foci in recipient cells (**Fig. 3d**). No Tau-FR foci were detected when donor cells expressed only soluble Tau-GFP (**suppl. figure 3c, d**). Increased numbers of recipient cells with visible Tau-FR foci were observed when cells were cocultured with VSV-G-expressing HEK Tau-GFP^AD^ donors (**Fig. 3d**). We also observed a significant increase in recipient cells with Tau-FR aggregates when HEK Tau-GFP^FTLD^, HEK Tau-GFP^PSP^ or HEK Tau-GFP^CBD^ were used as donors (**Fig. 3e**). VSV-G-expressing HEK Tau-GFP^agg^ cells also efficiently induced Tau-FR aggregation in human primary astrocytes expressing soluble Tau-FR (**suppl figure 3e, f**). Aggregate induction was dependent on the fusogenic activity of VSV-G, as coculture of recipients with donors expressing a non-fusogenic mutant resulted in strongly reduced Tau aggregate induction in recipients (**suppl. figure 3g, h**).

We next tested the effect of VSV-G expression on EV-mediated Tau aggregation. Sedimentation assays and Western blot analyses demonstrated that VSV-G pseudotyped EV fractions from HEK Tau-GFP^AD^ and Tau-GFP^CBD^ contained VSV-G, aggregated Tau-GFP and EV markers Flotillin1 and Hsp70/72 (**Fig. 3f, g**). Addition of VSV-G pseudotyped EV to recipient HEK Tau-FR^sol^ cells drastically increased Tau aggregate induction (**Fig. 3h-j**). Again, destruction of EV by sonication basically abolished aggregate induction, demonstrating that functional EV were required for transmission of seeding-competent Tau (**Fig. 3j**). VSV-G expression also increased the number of particles released by donors, but did not change the size distribution of EV (**Fig. 3k, l)**. VSV-G pseudotyped EV adjusted to comparable particle numbers also resulted in increased Tau aggregate induction (**Fig. 3m**). We conclude that decoration of EV with viral glycoprotein VSV-G mediates attachment and fusion of EV with the target cell, thereby efficiently transmitting seeding-competent Tau seeds.

### VSV-G-pseudotyped EV efficiently transmit scrapie prions to recipient cells

Transmissible spongiform encephalopathy (TSE) agents, the so far only *bona fide* mammalian prions, are composed of misfolded prion protein PrP ^6^. The conversion of cellular prion protein (PrP^C^), a protein tethered to the cell membrane by a glycosylphosphatidyl-anchor, into its infectious aggregated isoform PrP^Sc^, occurs on the cell surface or along the endocytic pathway ^37^. We tested the effect of VSV-G on the intercellular spreading of mouse-adapted scrapie in susceptible cell lines. It has been shown that N2a cells release prion infectivity associated with EV ^20, 38^. N2a cells persistently infected with TSE strain 22L (N2a^22L^) were transiently transfected with control plasmid or a plasmid coding for VSV-G (**Fig. 4a**). Murine fibroblast cell line L929 ^39^ and CAD5 cells ^40^, two cell lines highly permissive to TSE strain 22L, were exposed to VSV-G positive EV (**Fig. 4b**) and subsequently cultured in the absence of EV for more than 8 passages before testing for the formation of PrP^Sc^ (**Fig. 4a**). VSV-G pseudotyping strongly affected PrP^Sc^ accumulation in recipient cells, as determined by proteinase K treatment of cell lysates followed by Western blot analysis (**Fig. 4c**). VSV-G decorated EV also increased the number of cells containing PrP^Sc^ aggregates (**Fig. 4d, e**). We conclude that expression of viral ligand VSV-G drastically increases the capacity of donor cells to transmit both cytosolic and membrane-anchored proteopathic seeds to recipient cells.

**Fig. 4.**
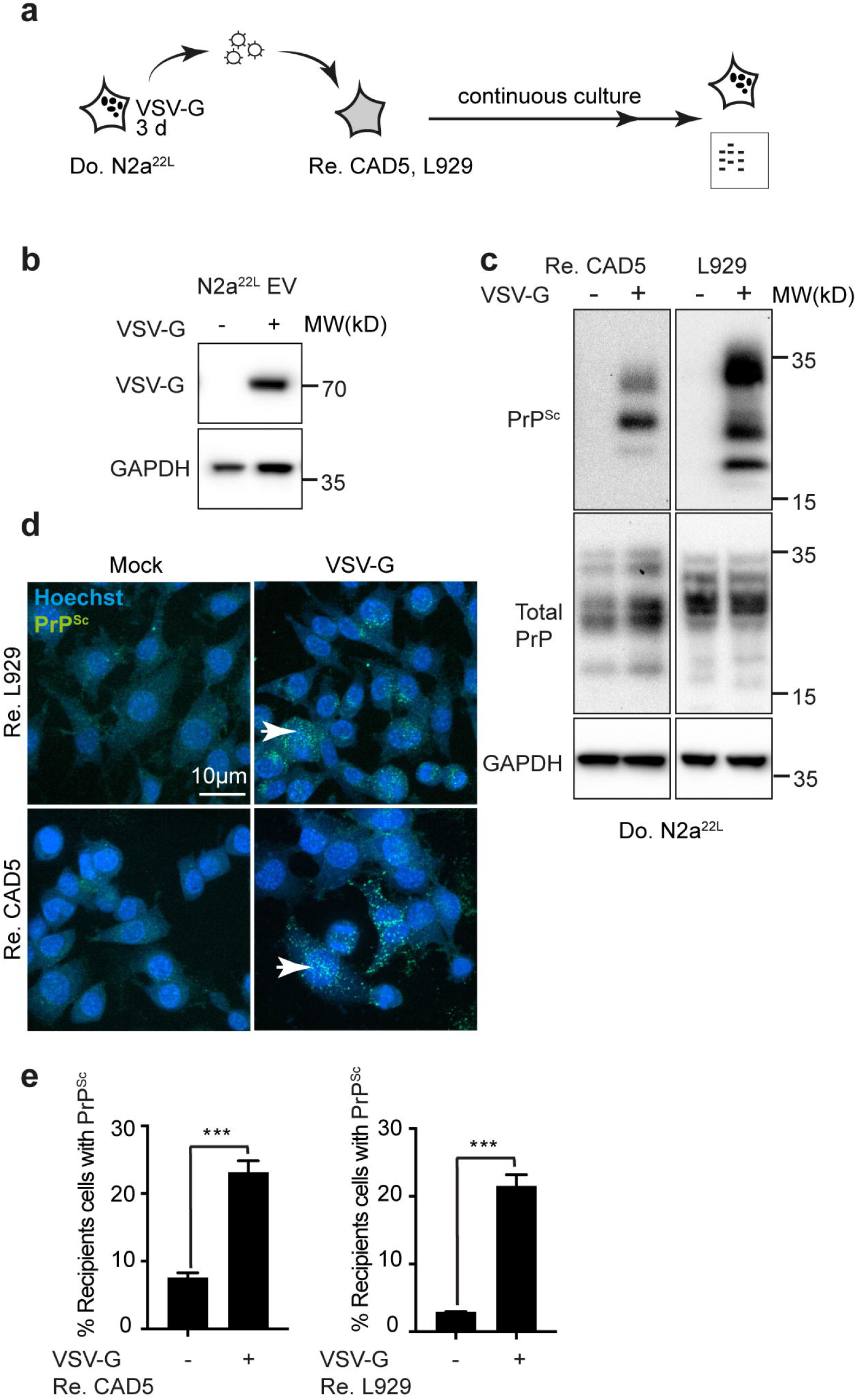
VSV-G increases spreading of transmissible spongiform encephalopathy agent. **a**. Experimental workflow. N2a cells persistently infected with mouse-adapted scrapie strain 22L (N2a^22L^) were transfected with empty vector or VSV-G coding vector. 3 d later, EV in conditioned medium were harvested and added to prion-permissive CAD5 or L929 cells. Subsequently, recipient cells were passaged 8 times to dilute out inoculum before accumulation of prions was monitored by immunofluorescence staining of Guandidinium hydrochlorid-(GdnHCL) unfolded PrP^Sc^ or proteinase K resistant PrP^Sc^ by Western blot. **b**. Detection of VSV-G and GAPDH in EV preparations from donor cells. **c**. Exposure of recipient cells to VSV-G pseudotyped EV from N2a^22L^ donors leads to efficient infection, as demonstrated by the presence of PK-resistant PrP^Sc^ in cells exposed to pseudotyped EV from scrapie-infected donors. GAPDH and total PrP were detected on a separate blot. **d**. Detection of PrP^Sc^ in recipient L929 and CAD cells exposed to EV from scrapie-infected donor cells. Eight passages post infection, recipient cells were fixed and protein was denatured using 6 M GdnHCL. Subsequently, PrP^Sc^ was detected using antibody 4H11. Arrowheads indicate PrP^Sc^ signal. **e**. Quantitative analyses of individual cells with PrP^Sc^ puncta following exposure to EV from transfected donor cells (as of d) (n=6, ***, *p*<0.001; unpaired student t test).

### Increased proteopathic seed spreading upon SARS-CoV-2 spike expression

Next, we tested if glycoproteins associated with human pathogenic viruses could contribute to protein aggregate spreading. SARS-CoV-2 is a novel Betacorona virus that has become a pandemic threat with millions of confirmed cases since its outbreak in December 2019 ^41^. SARS-CoV-2 binds to its target cells by interaction of its spike protein S with the human angiotensin-converting enzyme 2 receptor (ACE2) ^41-43^. Spike S protein is a large transmembrane protein that is cleaved by host proteases into two subunits responsible for receptor binding and fusion with the host cell membrane. We assessed if ectopic expression of spike S by donor cells modulates proteopathic seed spreading in our cellular models (**Fig. 5a**). To this end, donor cell populations propagating NM-HA or Tau-GFP aggregates were transfected with vector coding for SARS-CoV-2 spike S or pcDNA3.1(+) plasmid as Mock control. Both precursor protein and cleaved subunit S1 were identified in lysates of transfected cells, demonstrating that spike S was accurately processed by host proteases (**Fig. 5b**). HEK cells overexpressing ACE2 (**Fig. 5c**) or Vero cells highly susceptible to SARS-CoV-2 ^41^ were used as recipients (**Fig. 5a**). Coculture with aggregate-bearing HEK donors overexpressing spike S increased aggregate induction in recipients (**Fig. 5d, e**) and clearly depended on the expression of the viral ligand (**Fig. 5f-h**). Importantly, spike S also associated with the EV fraction secreted by donor cells (**Fig. 5i**). Isolated EV were also tested for their aggregate-inducing capacity in their respective target cells (**Fig. 5j**). Spike S expression did not affect EV secretion (**Fig. 5k**), but significantly increased numbers of recipient cells with induced aggregates (**Fig. 5l, m**). We conclude that glycoproteins from different viral genera facilitate the spreading of proteopathic seeds, suggesting that viral glycoproteins could generally contribute to the intercellular exchange of cellular components.

**Fig. 5.**
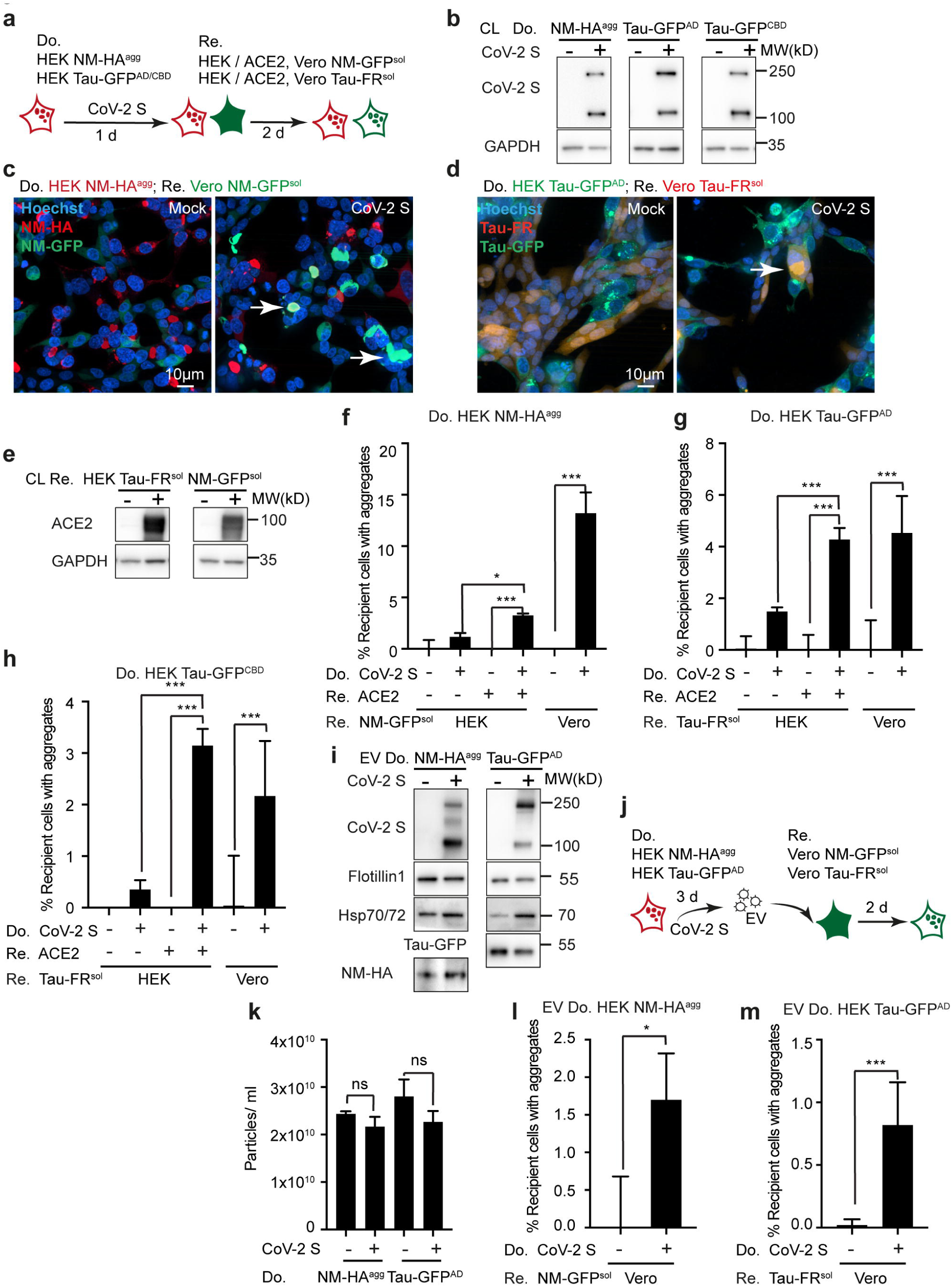
SARS-CoV-2 spike S protein affects spreading of proteopathic seeds. **a**. Schematic diagram of the experiment. HEK donor cells propagating NM-HA or Tau-GFP seeds were transfected with plasmid coding for CoV-2 spike S or Mock transfected. 1 d later, cells were cocultured with recipient HEK cells expressing the soluble aggregation-prone proteins and overexpressing/ not overexpressing human ACE2. Alternatively, donors were cultured with Vero cells stably expressing NM-GFP^sol^ or Tau-FR^sol^. Induction of protein aggregates was assessed 2 d later. **b**. Western blot demonstrating expression of Myc epitope-tagged CoV-2 spike S in donor HEK cells. GAPDH served as loading control. Note that spike S runs as two bands, representing the full-length precursor and the cleaved form. **c**. Representative images of cocultures of HEK NM-HA^agg^ donors and recipient Vero cells expressing soluble NM-GFP. Donor cells were either Mock transfected or transfected with vector coding for spike S protein. **d**. Representative images of cocultures of donor HEK Tau-GFP^AD^ cells with recipient Vero Tau-FR^sol^ cells. **e**. Western blot demonstrating ectopic expression of ACE2-Flag in recipient HEK cells. The blot was probed with antibodies against human ACE2. **f**.**-h**. Percentage of recipient HEK or Vero cells with induced NM-GFP^agg^, Tau-FR^AD^ and Tau-FR^CBD^ in the presence of donor cells with/ without spike S (n=6, *, *p*<0.1, ***, *p*<0.001; one-way ANOVA). HEK recipients were either Mock transfected or transfected with ACE2 coding vector. **i**. Presence of spike S on EV. EV isolated from transfected donor clones were subjected to Western blot analysis to assess the presence of S protein, NM-HA or Tau-GFP proteins and EV markers. **j**. Experimental design. HEK donor clones propagating proteopathic seeds were Mock transfected or transfected with plasmid coding for spike S. EV were subsequently added to recipient Vero cells. The percentage of recipients with induced aggregates was assessed 2 d later. **k**. Expression of spike S does not affect particle numbers. Statistical analysis was performed using one-way ANOVA (n=3). **l**., **m**. Percentage of EV-exposed recipient cells with induced NM-GFP^agg^ and Tau-FR^AD^ in the Vero HEK NM-GFP^sol^ or Tau-FR^sol^ recipient cells (n=3, *, *p*<0.1, ***, *p*<0.001; unpaired student t test).

## Discussion

Dissemination of protein aggregates between cells can occur by secretion of membrane-free naked aggregates, secretion as vesicular cargo or via direct cell-to-cell contact. As most disease-associated proteins are secreted in a membrane-free state, it is unclear how much cell-to-cell contact and EV-mediated transfer really contribute to protein aggregate spreading ^10^. We reasoned that proteopathic seed transmission involving the latter two pathways is at least partially controlled at the cell entry step. Here we demonstrate that decoration of EV with viral glycoproteins VSV-G or SARS-CoV-2 spike S, which mediate receptor interaction and subsequent merger of opposing membranes, increases protein aggregate induction in recipient cells. Thus, decoration of EV with ligands that mediate effective interaction and fusion with target cells turns them into potent delivery vehicles for proteopathic seeds. The fact that intercellular dissemination of three independent proteopathic seeds could be strongly increased by efficient receptor-ligand interactions clearly shows that mechanisms of intercellular protein aggregate transfer are overlapping. Moreover, the effect of viral glycoproteins on seed transmission suggests that certain viral infections could contribute to the dissemination of proteopathic seeds and ultimately modulate progression of protein misfolding diseases.

Recent findings that disease-associated proteins such as Tau and α-synculein are predominately secreted in a non-membrane bound state ^17, 19, 22, 44^ have focused research on uptake mechanisms of free oligomers or fibrils, thereby uncovering heparan sulfate proteoglycans ^44, 45^ and Lrp1 ^46^ as receptors. Still, mechanisms of seed dissemination are not mutually exclusive. The fact that mechanical destruction of EV basically abolished aggregate induction in recipients suggests that, in our experimental setting, vesicle-mediated transfer is much more efficient than uptake of naked seeds from ruptured EV. Importantly, we previously showed that oligomeric species of NM aggregates packaged in EV were not affected by sonication ^33^. Thus, EV equipped with suitable ligands represent highly effective vehicles for transfer of seeding-competent cargo. By contrast, insufficient receptor-ligand interactions constitute barriers to EV-mediated proteopathic seed spreading that can obscure the actual seeding capacity of packaged cargo.

Previous studies have demonstrated that VSV-G-pseudotyped EV share exosome markers and fall in the size range of exosomes that originate from intraluminal budding of vesicles into late endosomes ^28^. This is consistent with our findings, demonstrating a consistent EV size of approx. 100 nm unaffected by viral glycoprotein expression. Based on previous evidence, neither size nor marker proteins can specifically discriminate exosomes from plasma membrane-shed vesicles, making such discriminations impossible ^16^. Noteworthy, proteopathic seeds have been found secreted both as small exosome-like vesicles and large membrane-shed vesicles, arguing that aggregates composed of the same disease-associated protein can be released from different cellular sites ^19, 21, 47, 48^.

Expression of viral envelope protein VSV-G not only increased cell-free transmission of proteinaceous seeds by EV, but also promoted aggregate induction in recipient cells in coculture. Intriguingly, transmission routes of proteopathic seeds resemble those used for viral spreading. In addition to dissemination via released mature viral particles, viruses have evolved sophisticated strategies to exploit intercellular communication for non-classical dissemination. Mechanisms of cell-to-cell viral spread include fusion of cellular plasma membranes, intercellular membranous connections such as cytonemes or TNTs, or virus-induced cell interfaces reminiscent of tight junctions or synapses ^49^. Mature viral particles packaged into vesicles can also be transported through the lumen of TNTs ^50^. Retroviruses and some herpesviridae exploit cytonemes emanating from infected cells to transfer infectious virus surfing on the outer surface to connecting cells ^51^. Viral infection through these direct cell-to-cell contacts can be 2-3 orders of magnitude more efficient than by released viruses. Enveloped and non-enveloped viruses such as hepatitis A virus efficiently exploit EV for non-lytic cellular egress and subsequent infection ^52^. HIV glycoprotein and its receptor concentrate at filopodial tips of donor and recipient cells, thereby likely driving tip fusion and establishment of cytonemes for efficient cell-to-cell infection ^51^.

Microbial brain infections have long been suspected to play a role in pathogenesis of neurodegenerative diseases ^53^. For example, co-infection of fibroblasts with scrapie and retroviral Moloney leukemia virus resulted in the increased secretion of prions associated with EV, an effect attributed to the expression of Gag capsid protein ^54^. Gag expression also increased persistence of prion infection in an epithelial cell line infected with chronic wasting disease prions ^55^. It is tempting to speculate that viral fusogens, expressed during acute or latent human infections, could affect spreading of proteopathic seeds. VSV is a rhabdovirus infecting ungulates that occasionally causes zoonotic flu-like infections in humans, and thus does not play a role in proteopathic seed spreading in neurodegenerative diseases. CoV-2 in humans primarily manifests as a respiratory illness, but neurological symptoms are present in 25 % of acute cases and can be linked to direct involvement of the central nervous system ^56^. Neuroinvasion, as evident by the presence of SARS CoV-2 viral RNA in the CNS ^57, 58^, can occur through the olfactory transmucosal route ^59^. Although brain cell infection is still under debate, both astrocytes and neurons express ACE2 and thus may represent SARS CoV-2 target cells ^60, 61^. COVID-19 infections could increase the risk for developing neurological or neurodegenerative diseases later in life due to direct or indirect effects on the CNS, such as viral encephalitis, systemic organ failure, cerebrovascular complications or inflammation ^56^. However, if and how COVID-19 infection affects disease pathogenesis in ND remains to be established. Importantly, several neurotrophic viruses causing lifelong persistent infections such as herpesviridae are upregulated in the CNS of ND patients ^62, 63^. The expression of viral glycoproteins could thus increase cell-to-cell spreading of protein aggregates and also expand the host cell tropism. Future research is required to clarify the role of viral infections and specific viruses in the prion-like progression of protein aggregation in neurodegenerative and other protein misfolding diseases.

## Methods

### Human brain samples

Frozen postmortem brain tissue samples from neuropathologically confirmed cases of AD, FTLD-Tau, CBD, PSP and controls were provided by the Brain Bank associated with the University Hospital and DZNE Tübingen. All material was collected from donors for or from whom a written informed consent for a brain autopsy and the use of the material and clinical information for research purposes had been obtained. All samples are listed in Supplementary Table 1.

### Ethics statement

Ethical approval for usage of human samples for the current study was obtained from ‘Medizinische Fakultät Ethik-Kommission, Rheinische Friedrich-Wilhelms-Universität, project no. 236/18(2018)’.

### Molecular cloning

For lentiviral constructs Tau-GFP /-Fusion Red (FR), human four repeat domain (4R) Tau (amino acids 243 to 375) with mutations P301L and V337M was fused to GFP or FR (Evrogen) with an 18-amino acid flexible linker (EFCSRRYRGPGIHRSPTA), as described previously (thereafter termed Tau-GFP, Tau-FR) ^64^. Coding regions were cloned into the lentiviral vector pRRL.sin.PPT.hCMV.Wpre via BamHI and SalI ^32^. For the generation of non-fusogenic VSV-G mutants ^65^, mutations were inserted into the open reading frame of Vesicular stomatitis virus glycoprotein VSV-G using the Q5 SDM kit (NEB). Myc epitope-tagged SARS-CoV-2 (2019-nCoV) spike S cDNA (VG40589-CM) and Flag epitope-tagged human ACE2 cDNA (HG10108-NF) plasmids were purchased from Sino Biological.

### Cell lines

N2a, L929, CAD5 and HEK 293T cells were cultured in Opti-MEM (Gibco) supplemented with glutamine, 10 % (v/v) fetal bovine serum (FCS) (PAN-Biotech GmbH) and antibiotics. Human primary astrocytes (ScienCell) were cultivated as recommended by ScienCell. Vero cells were purchased from CLS (Cell lines service) and cultivated as recommended. All cells were incubated at 37°C and 5% CO_2_. The total numbers of viable cells and the viability of cells were determined using the Vi-VELL™XR Cell Viability Analyzer (Beckman Coulter). Transfection of cells were performed either with lipofectamine2000 or TransIT-2020 / X2 (Mirus) reagents as recommended by the manufacturers.

### Production and transduction with lentiviral particles

HEK293T cells were cotransfected with plasmids pRSV-Rev, pMD2.VSV-G, pMDl.g/pRRE, and pRRl.sin.PPT.hCMV.Wpre containing Tau-GFP/FR using lipofectamine2000. Supernatants were harvested and concentrated with PEG method according to published protocols ^66^. Cell lines and primary neurons were transduced with lentivirus, and stable cell clones expressing Tau-GFP/FR were selected following limiting dilution cloning ^31^.

### Extracellular vesicle isolation

To prepare exosome-depleted medium, FCS was ultracentrifuged at 100,000 x g, 4 °C for 20 h. Medium supplemented with the exosome-depleted FCS and antibiotics was subsequently filtered through a 0.22 and a 0.1 µM filter-sterilization device (Millipore). For EV isolation, 2-4×10^6^ cells were seeded in a T175 flask in 35 ml exosome-depleted medium to be confluent after 3 days. Cells and cell debris were pelleted by differential centrifugation (300 x g, 10 min; 2,000 x g, 20 min; 16,000 x g, 30 min). The remaining supernatant (conditioned medium) was subjected to ultracentrifugation (UC) (100,000 x g 1 h) using rotors Ti45 or SW32Ti (Beckman Coulter). The pellet was rinsed with PBS and spun again using rotor SW55Ti (100,000 x g 1 h).

### Aggregate induction assay

Recipient cells were cultured on CellCarrier-96 or 384 black microplate (PerkinElmer) at appropriate cell numbers for 1 h, and then treated with 5-10 µl of EV. For coculture, a total of 10^4^ cells/ per well of recipient and donor cells was plated at different ratios depending on their population doubling times. After additional 1 day for NM expressing and 2 days for Tau expressing cultures, cells were fixed in 4 % paraformaldehyde and nuclei were counterstained with Hoechst. Cells were imaged with the automated confocal microscope Cellvoyager CV6000 (Yokogawa Inc.) using a 20 x or 40 x objective. Maximum intensity projections were generated from Z-stacks. Images from 16 fields per well were taken. On average, a total of 3-4×10^4^ cells per well and at least 3 wells per treatment were analyzed.

### Determination of extracellular vesicles size and number

ZetaView PMX 110-SZ-488 Nano Particle Tracking Analyzer (Particle Metrix GmbH) was used to determine the size and number of isolated extracellular vesicles. The instrument captures the movement of extracellular particles by utilizing a laser scattering microscope combined with a video camera. For each measurement the video data is calculated by the instrument and results in a velocity and size distribution of the particles. For nanoparticle tracking analysis, the Brownian motion of the vesicles from each sample was followed at 22° C with properly adjusted equal shutter and gain. At least six individual measurements of 11 positions within the measurement cell and around 2200 traced particles in each measurement were detected for each sample.

### Sedimentation assay for Tau

Sedimentation assay was performed as described previously ^36^. Briefly, cleared cell lysates with 100 µg total protein was centrifuged at 100,000 x g for 1 h. Pellet was washed with 1.5 ml PBS and centrifuged at 100,000 x g for 30 min. Supernatant fractions were precipitated with 4 x methanol at −20° C overnight, and protein was pelleted at 2,120 x g for 25 min at 4° C (soluble fraction). The pellet (insoluble fraction) and 1/3 of the soluble fraction dissolved in RIPA buffer with 4 % SDS were loaded for western blot analysis.

### Pronase digestion for Tau

Pronase digestion experiment was performed as described previously ^36^. Briefly, 18 µl cleared cell lysate or brain homogenates (20-100 µg based on Tau aggregate content) were incubated with 2 µl 1 mg / ml pronase (Roche) at 37° C for one hour. Afterwards, reactions were boiled with 3 x sample buffer, and Tau was detected by Western blot as described below.

### Brain homogenate preparation and clarification

Frozen human brain samples were homogenized in lysis buffer (for protein analysis) via Precellys® 24 (Bertin Instruments) with 1.4 mm ceramic beads at 4° C for 4 cycles 5500 rpm 20 s. To prepare 10 % (w/v) clear brain homogenate for aggregate induction, crude homogenates were centrifuged at 872 x g for 5 min at 4° C, and then the supernatants were sonicated with 50 % power for 6min. These homogenates were frozen at −80° C until use. For protein analysis, cleared supernatants were prepared by centrifugation of the crude homogenates at 15,000 x g for 15 min.

### Tau aggregate induction by brain homogenate and liposomes

To induce Tau aggregation in HEK Tau-GFP^sol^ clone with brain homogenates from different tauopathy patients, cells were plated on 6-well plates at 1×10^6^ cells/ well in 2 ml complete medium one day before. Next day, 200 µl 10 % brain homogenates and 4 µl lipofectamine2000 were incubated for 20 min and added to recipient cells to have final 1 % brain homogenates on cells. After 3 days, cells were split and further expanded for limited dilution clone selection as previous described.

### PK treatment for PrP^Sc^

Cell pellet collected from one well of 6 well plate was lysed in 1ml lysis buffer. 900 µl of lysates were digested with 20 µg/ ml proteinase K (PK) at 37° C for 30 min for PrP^Sc^ detection. Proteolysis was terminated by addition of 0.5 mM Pefabloc. Rest of the 100 µl lysates for PrP^C^ detection and the digested samples were precipitated with methanol and analyzed by Western blot using anti-PrP antibody 4H11.

### Western blotting

For Western blot analysis, protein concentrations were measured by Quick Start™ Bradford Protein assay (Bio-Rad) and proteins were separated on NuPAGE®Novex® 4-12 % Bis-Tris Protein Gels (Life Technologies) followed by transfer onto a PVDF membrane (GE Healthcare) in a wet blotting chamber. Western blot analysis was performed using mouse anti-Alix (1:1000; BD Bioscience); rat anti-HA 3F10 (1:1000; Roche); mouse anti-GAPDH 6C5 (1:5000; Abcam); mouse anti-Hsc/Hsp70 N27F3-4 (1:1000; ENZO); mouse anti-VSV-G A5977 (Sigma); rabbit anti-Tau ab64193 (Abcam); rabbit anti-Flotillin 1 ab133497 (Abcam); mouse anti-SARS-CoV-2-spike S GTX632604 (GeneTex); rabbit anti-hACE2 ab15348 (Abcam). The membrane was incubated with Pierce™ ECL Western Blotting Substrate (Thermo Fisher Scientific) according to the manufacturer’s recommendations.

### Automated image analysis

The image analysis was performed using the Cellvoyager Analysis support software. An image analysis routine was developed for single cell segmentation and aggregate identification (Yokogawa Inc.) The total number of cells was determined based on the Hoechst signal, and recipient cells were detected by their GFP / FR signal. Green aggregates were identified via morphology and intensity characteristics. The percentage of recipient cells with aggregated NM-GFP or Tau-FR/ Tau-GFP was calculated as the number of aggregate-positive cells per total recipient cells set to 100 %. False positive induced recipient cells were detected due to the heterogeneity in GFP / FR expression of individual cells. The mean percentage of false positives determined in control recipient cells was subtracted from all samples. Of note, negative values were sometimes obtained when no induction was observed. For data presentation, the minimum range of Y Axis was set to 0.

### Immunofluorescence staining and confocal microscopy analysis

Cells were fixed in 4 % paraformaldehyde and permeabilized in 0.1 % Triton X-100. HA staining was performed with Alexa Fluor 647-conjugated anti-HA antibody (MBL M180-A647). For PrP^Sc^ staining, proteins were denatured in 6 M guanidine hydrochloride for 10 min at RT to reduce the PrP^C^ signal ^67^. Cells were rinsed with PBS, blocked in 0.2 % gelatine and incubated for 2 h with 4H11 antibody diluted 1:10 in blocking solution. After three washing steps in PBS, cells were incubated for 1 h with Alexa Fluor 488-conjugated secondary antibody and nuclei were counterstained for 15 min with 4 µg/ml Hoechst 33342 (Molecular Probes). 96 and 384 well plates were scanned with Cellvoyager CV6000 (Yokogawa Inc.). Confocal laser scanning microscopy was performed on a Zeiss LSM 800 laser-scanning microscope with Airyscan (Carl Zeiss).

### Statistical analysis

All analyses were performed using the Prism 6.0 (GraphPad Software v.7.0c). Statistical inter-group comparisons were performed using the one-way ANOVA with a Bonferroni post-test or Student’s t test. p values smaller than 0.1 were considered significant. All experiments were performed in triplicates or sextuplicates and repeated at least two times. Error bars represent the standard deviation (SD).

## Acknowledgements

We are grateful to Birgit Kurkowsky of Laboratory Automation Technologies, DZNE Bonn, for technical support regarding automated imaging acquisition and analysis. We thank Corinne Lasmezas, Department of Immunology and Microbiology, Scripps Research Institute Florida, for providing prion-susceptible CAD5 cells. Funding was obtained from the Helmholtz Portfolio Wirkstoffforschung and from the Deutsche Forschungsgemeinschaft (DFG, German Research Foundation) under Germany’s Excellence Strategy within the framework of the Munich Cluster for Systems Neurology (EXC 2145 SyNergy– ID 390857198).

## Author contributions

S.L., and I.V. designed research; S.L., A.H., A.H. and O.B. performed research; M.N. contributed human samples; S.L., S.F.L., S.M., P.D. and I.V. analyzed data; and S.L. and I.V. wrote the paper.

