## Supplemental Fig Legend for "Highly efficient intercellular spreading of protein misfolding mediated by viral ligand - receptor interactions"

**Suppl. figure 1.** Representative images of coculture and EV experiments. **a.** Experimental workflow for coculture. Two donor cell clones, HEK NM-HA$^{agg}$ (red) and N2a NM-HA$^{agg}$ (red), with poor aggregate inducing activity were cocultured with recipient HEK or N2a cells expressing NM-GFP$^{sol}$ (green). Aggregate induction was monitored 24 h later. **b.** Experimental workflow for EV-mediated aggregate induction assay using the same donor and recipient cell clones as in (a). EV from donor clones were incubated with either N2a or HEK cells expressing soluble NM-GFP for 1 d. **c. d.** Recipient N2a NM-GFP$^{sol}$ cells cocultured (left panel) or exposed to donor EV (right panel). Donor cells were either N2a NM-HA$^{agg}$ (c) or HEK NM-HA$^{agg}$ (d). **e., f.** Recipient HEK NM-GFP$^{sol}$ cells cocultured with donor cells (left panel) or exposed to donor EV (right panel). Donor cells were either N2a NM-HA$^{agg}$ (e) or HEK NM-HA$^{agg}$ (f). NM-HA was stained using anti-HA antibodies.

**Suppl. figure 2. a.** Cocultures of donor N2a NM-HA$^{agg}$ cells Mock or VSV-G transfected with either N2a or HEK cells expressing soluble NM-GFP. NM-HA was stained using anti-HA antibodies. **b.** Occasional syncytia formation indicates fusogenic activity of VSV-G. Arrowheads indicate multinucleated cells with protein aggregates positive for HA and GFP. **c.** Experimental design controls. Control donor N2a NM-HA$^{sol}$ cells were transfected with VSV-G plasmid or Mock transfected. The following day, donor cells were cocultured with recipient N2a or HEK cells expressing NM-GFP$^{sol}$. **d.** Representative images of cocultures. Arrowheads mark occasional multinucleated cells.

**Suppl. figure 3. a.** Different tauopathy patient brain homogenates contain Tau aggregates with distinct pronase-resistant patterns. Tau was detected using ab64193. **b.** Tau-GFP in HEK Tau-GFP$^{agg}$ cells is pronase resistant. Individual HEK clones propagating Tau-GFP aggregates and control HEK Tau-GFP$^{sol}$ cells were lysed and subjected to pronase treatment. Tau-GFP was detected using anti-Tau antibody (ab64193). **c.** Experimental design controls. Control donor HEK Tau-GFP$^{sol}$ was transfected with VSV-G plasmid or Mock transfected. The following day donor cells were cocultured with recipient HEK cells expressing Tau-FR$^{sol}$. **d.** Representative images of cocultures. Arrowheads mark occasional multinucleated cells. **e.** Donor clone HEK Tau-GFP$^{AD}$ transfected or not with VSV-G plasmid was cocultured with human astrocytes ectopically expressing Tau-FR$^{sol}$. **f.** Transfected donor clones HEK Tau-GFP$^{AD}$ and HEK Tau-GFP$^{CBD}$ were cocultured with human astrocytes expressing Tau-FR$^{sol}$. Shown is the percentage of human astrocytes expressing Tau-FR$^{sol}$ with induced Tau-FR aggregates 2 days post coculture. Statistical analysis was performed using student´s unpaired t-test (n=6, ***, $p<0.001$). **g.** Expression of fusogenic (WT) and non-fusogenic VSV-G mutant upon transfection of HEK Tau-GFP$^{AD}$ and HEK Tau-GFP$^{CBD}$ donor cells. GAPDH served as loading control. **h.** Percentage of recipient cells cocultured with donors that contain Tau-FR$^{agg}$ and were transfected with Mock vector, VSV-G or its mutant (n=6, ***, $p<0.001$; one-way ANOVA).
