## Supplemental Table 1 for "Highly efficient intercellular spreading of protein misfolding mediated by viral ligand - receptor interactions"

**Supplementary Table 1**

| Patient No. | Age at death | Sex | NP-Diagnosis | ABC score* | Brain region |
|---|---|---|---|---|---|
| **1** | 65 | M | AD | A3, B3, C3 | frontal cortex |
| **2** | 63 | F | CBD | A2, Bx, C1 | frontal cortex |
| **3** | 77 | M | PSP | A1, B1, C1 | pons |
| **4** | 56 | M | FTLD-tau (*MAPT* IVS10+3 G>A) | A0, Bx, C0 | frontal cortex |
| **5** | 79 | F | Control | A1, B1, C0 | frontal cortex |

*ABC score according to the National Institute of Aging-Alzheimer's association guidelines [1].

1. Montine, T.J. *et al.* National Institute on Aging-Alzheimer's Association guidelines for the neuropathologic assessment of Alzheimer's disease: a practical approach. *Acta Neuropathol* **123**, 1-11 (2012).
