## Supplementary figures and images for "Highly efficient intercellular spreading of protein misfolding mediated by viral ligand - receptor interactions"

### Supplemental Fig. 1

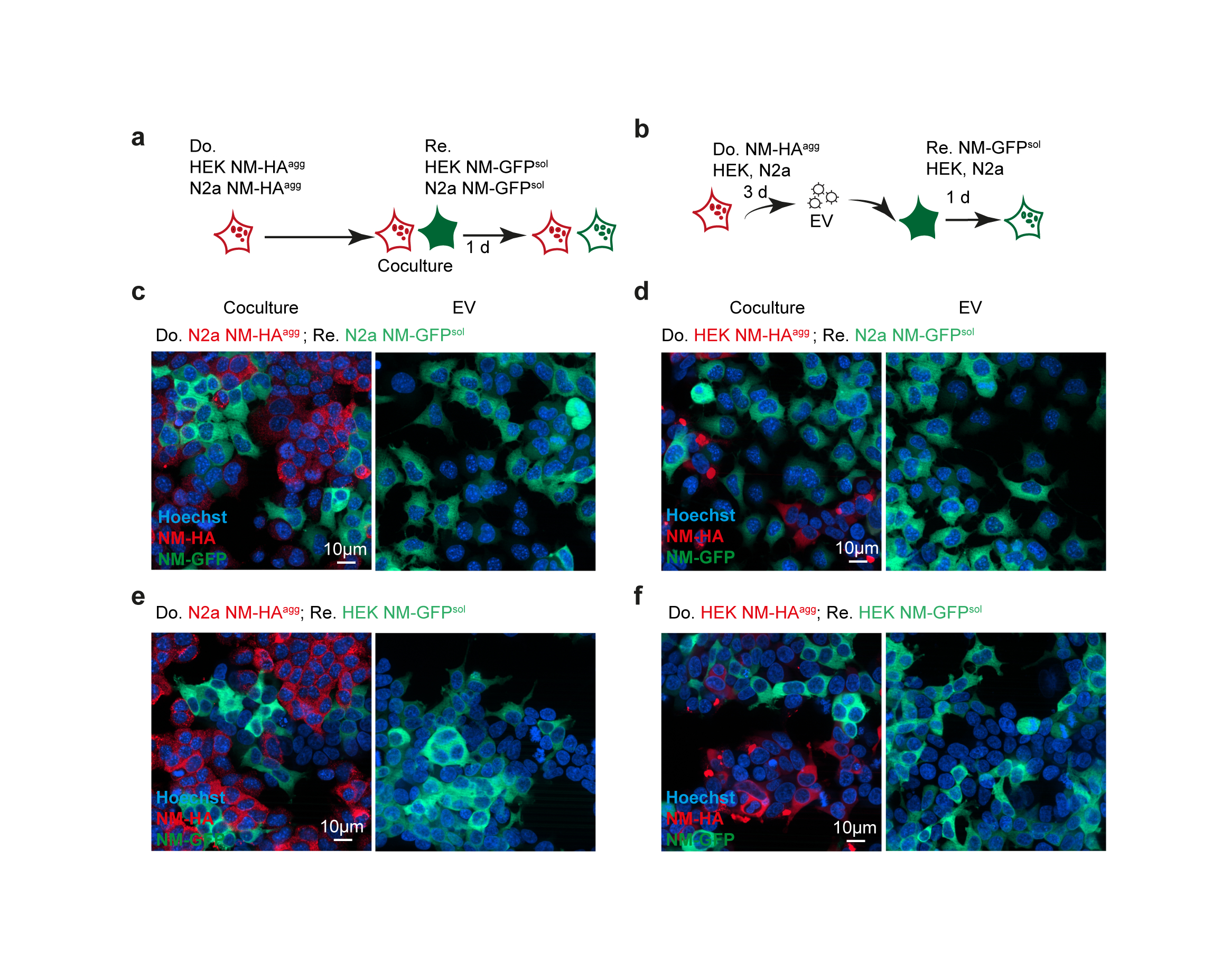

### Supplemental Fig. 2

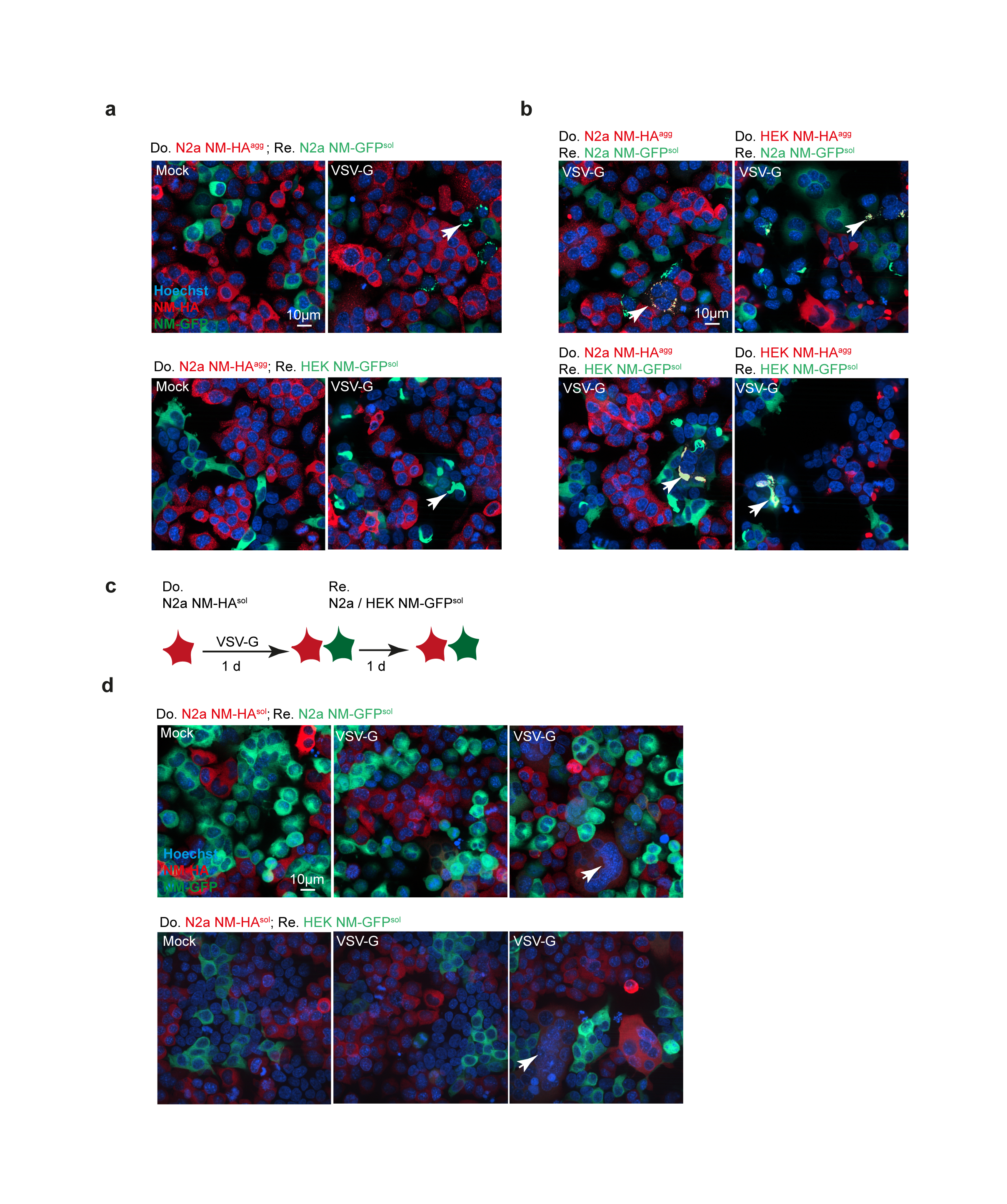

### Supplemental Fig. 3

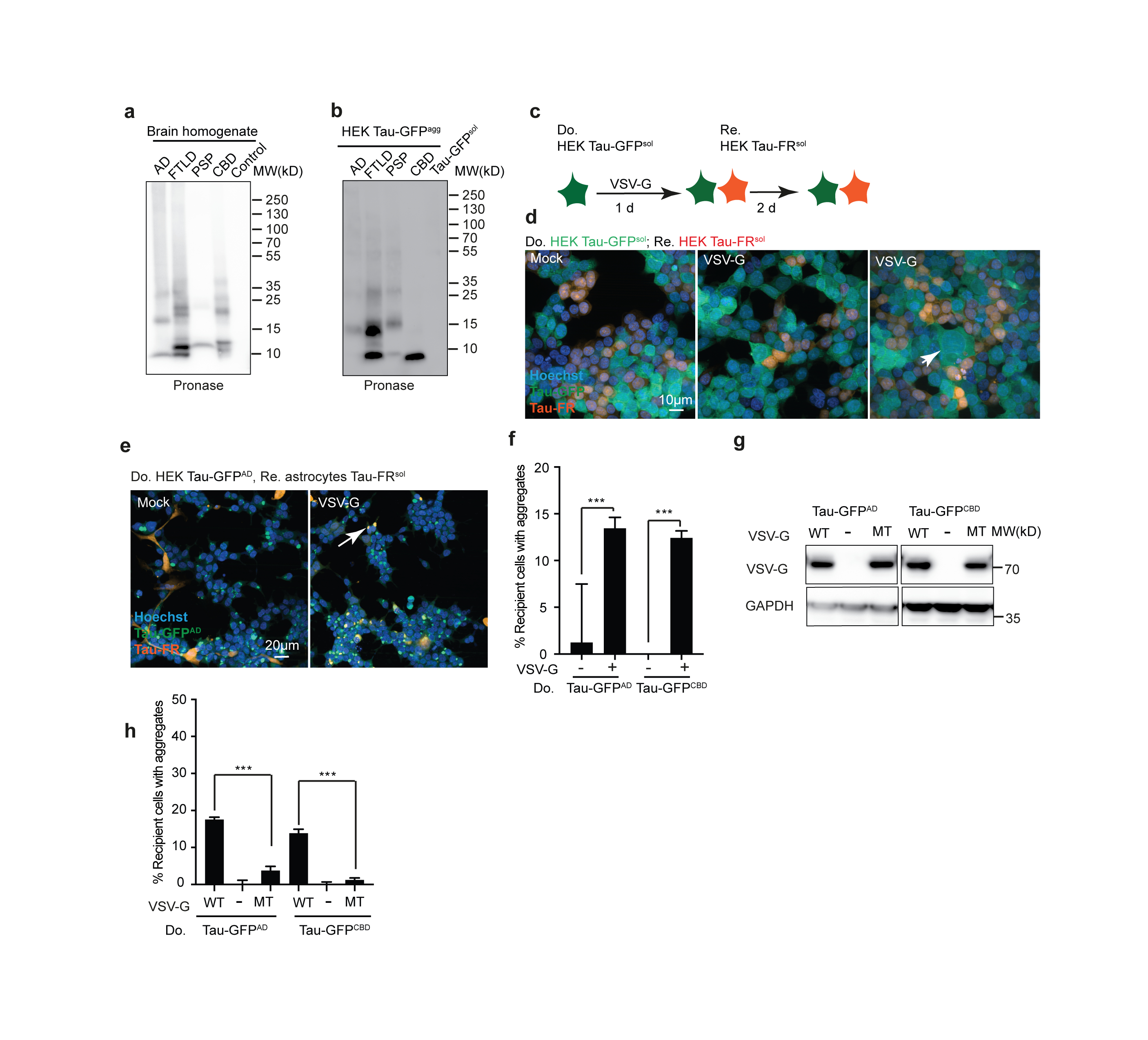
